# Massive-scale biological activity-based modeling identifies novel antiviral leads against SARS-CoV-2

**DOI:** 10.1101/2020.07.27.223578

**Authors:** Ruili Huang, Miao Xu, Hu Zhu, Catherine Z. Chen, Emily M. Lee, Shihua He, Khalida Shamim, Danielle Bougie, Wenwei Huang, Mathew D. Hall, Donald Lo, Anton Simeonov, Christopher P. Austin, Xiangguo Qiu, Hengli Tang, Wei Zheng

**Author notes:** Address correspondence and reprint requests to Ruili Huang, Ph.D., 9800 Medical Center Drive, DPI/NCATS, National Institutes of Health, Rockville, MD 20850.

## Abstract

The recent global pandemic caused by the new coronavirus SARS-CoV-2 presents an urgent need for new therapeutic candidates. While the importance of traditional *in silico* approaches such as QSAR in such efforts in unquestionable, these models fundamentally rely on structural similarity to infer biological activity and are thus prone to becoming trapped in the very nearby chemical spaces of already known ligands. For novel and unprecedented threats such as COVID-19 much faster and efficient paradigms must be devised to accelerate the identification of new chemical classes for rapid drug development. Here we report the development of a new biological activity-based modeling (BABM) approach that builds on the hypothesis that compounds with similar activity patterns tend to share similar targets or mechanisms of action. In BABM, compound activity profiles established on massive scale across multiple assays are used as signatures to predict compound activity in a new assay or against a new target. We first trained and validated this approach by identifying new antiviral lead candidates for Zika and Ebola based on data from ~0.5 million compounds screened against ~2,000 assays. BABM models were then applied to predict ~300 compounds not previously reported to have activity for SARS-CoV-2, which were then tested in a live virus assay with high (>30%) hit rates. The most potent compounds showed antiviral activities in the nanomolar range. These potent confirmed compounds have the potential to be further developed in novel chemical space into new anti-SARS-CoV-2 therapies. These results demonstrate unprecedented ability using BABM to predict novel structures as chemical leads significantly beyond traditional methods, and its application in rapid drug discovery response in a global public health crisis.

## Introduction

The early-stage drug discovery process relies on target identification, assay development, and high throughput screening (HTS) to identify lead compounds for chemical optimization and further preclinical development. Traditional HTS campaigns are often limited to 1-2 million compounds due to the high costs and operational bottle neck that limited the chance for lead identification.^1,2^ While it has been estimated that some 10^8^ drug-like compounds have been synthesized at least once, and that there are ~10^60^ drug-like molecules in chemical space, neither number is practical for HTS. However, recent advances in computational technologies have made it possible to virtually screen millions of compounds for potential biological activity.^2^ Existing virtual screening (VS) methods can be grouped into two broad categories: ligand-based VS and target structure-based VS. Ligand-based VS relies on the analysis of quantitative structure-activity relationship (QSAR) of existing experimental data obtained from either mechanistic or phenotypic compound screens, while target structure-based VS uses known protein structure information to predict the binding potential of small molecule compounds to protein targets. Both methods depend on chemical structure information to make predictions while the target-based approach in addition requires the availability of detailed target protein information. These severe dependencies have tended to limit the applicability of such methods to querying only in the close structural vicinity of already known ligand structures and drug targets.

A critical advance that enabled the development of activity rather than structural paradigm described here was the large-scale application of quantitative HTS (qHTS)^3^. Every compound in an qHTS campaign is tested in a broad concentration response format as opposed to in traditional HTS where compounds are tested only at single concentrations. The high-quality data from qHTS are thus substantially richer for use in computational modeling to predict activities of large compound libraries against new assays or new drug targets. In the past 15 years, our in-house collections of over half a million compounds have been screened in a wide spectrum of biological assays in qHTS format, which have been made publicly available,^4^ resulting in compound activity profiles that enabled the development of a new, biological activity-based modeling (BABM) approach complementary to traditional structure-based approaches.

Unlike traditional QSAR approaches (part of the ligand-based VS category),^5,6^ where similarity in chemical structure is used to infer biological activity, BABM builds on the hypothesis that compounds show similar activity patterns tend to share similar targets or mechanisms of action.^7,8^ In this approach, each assay is treated as an independent descriptor. Analogous to structure descriptors, where the presence and absence of certain structure features or properties are used to represent a compound, the presence and absence of activities against a panel of assays form the activity profile or signature of a compound. If extracted from across multiple screening campaigns each at massive scale, such ***activity signatures*** can then be applied to infer compound activity in a completely new assay or against a completely new target.

Analogous to a structural fingerprint, where the identity of each structural feature encoding the fingerprint does not need to be known, only the pattern of compound activity across assays is critical in BABM to make predictions whereas the identity of the individual assays forming the pattern is not important. A fundamental difference compared to traditional QSAR modeling is thus that BABM does not require any chemical structure information to make predictions, such that its application domain is not limited to small molecules with well-defined structures. In fact, BABM can be applied to any substances with available biological profiling, including macromolecules and mixtures (e.g. natural products).

Of particular note is that compounds showing similar activities do not necessarily share similar structures. Thus the BABM approach has no intrinsic limitations in discovering new chemical scaffolds in contrast to traditional QSAR methods, and can be used to virtually screen new compound libraries for which no activity data is available. These new scaffolds can then serve as starting points for lead identification efforts and be used to construct new QSAR models for lead optimization.

Starting in late fall 2019, a highly contagious disease, coronavirus disease 2019 (COVID-19), caused by a new coronavirus, severe acute respiratory syndrome coronavirus 2 (SARS-CoV-2), emerged and quickly spread across the globe.^9^ As of June 2020, over 8.2 million people have been infected with over 440,000 fatalities (https://covid19.who.int/). Despite some promising results from repurposed drug candidates that were shown to shorten the course of the disease (remdesivir^10,11^) or reduce mortality (dexamethasone^12,13^), other efforts have been disappointing or controversial (e.g., Lopinavir-Ritonavir^14^, hydroxychloroquine (HCQ)/ chloroquine (CQ)^15,16^), and an effective treatment has yet to be found for this disease.

This presented an urgent need for new methods that can quickly and systematically screen large compound libraries for new drug candidates. In this context, we first applied BABM to generate prediction models for two infectious diseases, Zika and Ebola, to test the robustness of BABM and its applicability to different assay and data types, and to benchmark against traditional QSAR methods,. Zika virus (ZIKV) belongs to the family of flaviviruses that have caused global epidemics resulting in human fatality.^17–19^ The flavivirus non-structural protein NS1 has been found to interact with a number of host molecules, and play many diverse roles in viral replication and virion maturation.^20,21^ The large outbreaks of Ebola virus (EBOV) disease in recent years represents an ongoing global crisis.^22^ qHTS campaigns using *in vitro* EBOV assays have identified compounds that blocked virus entry into cells and suppressed viral replication, but the identified active compounds exhibited low anti-EBOV potencies.^23,24^

The BABM model identified actives that were experimentally verified with high confirmation rates (~50-80%). The approach was then applied to build prediction models for SARS-CoV-2. Among our in-house libraries, the NCATS Pharmaceutical Collection (NPC)^25^ and the Library of Pharmacologically Active Compounds (LOPAC) have been screened in nearly every one of our ~2,000 assays providing the most comprehensive set of activity profiles that comprise an ideal training dataset for machine learning models. To build prediction models for these disease targets, we selected training data that included both qHTS assay data (SARS-CoV-2 and ZIKV NS1^26^) and data collected from published literature (SARS-CoV-2 and EBOV^24^). These models, mostly trained on the qHTS activity profiles of the NPC and LOPAC library compounds, were applied to predict the activity of all ~0.5 million compounds in our in-house library. Models were constructed using BABM and the performances were compared with those of traditional QSAR models as well as a combination of both activity and structural features. A little over 300 compounds identified by the BABM models as potential anti-SARS-CoV-2 leads were then tested in a live virus assay with ~100 confirmed (>30%), validating the utility and accuracy of the BABM approach. Some of the experimentally confirmed lead compounds may have the potential to be further developed into new antiviral therapies.

## Materials and Methods

### SARS-CoV-2 cytopathic effect (CPE) assay

Vero-E6 cells previously selected for high ACE2 expression ^27^ (grown in EMEM, 10% FBS, and 1% Penicillin/Streptomycin) were cultured in T175 flasks and passaged at 95% confluency. Cells were washed once with PBS and dissociated from the flask using TrypLE. Cells were counted prior to seeding. A CPE assay previously used to measure antiviral effects against SARS-CoV ^28^ was adapted for performance in 384 well plates to measure CPE of SARS CoV-2 with the following modifications. Cells, harvested and suspended at 160,000 cells/ml in MEM/1% PSG/1% HEPES supplemented 2% HI FBS, were batch inoculated with SARS CoV-2 (USA_WA1/2020) at M.O.I. of approximately 0.002 which resulted in approximately 5% cell viability 72 h post infection. Compound solutions in DMSO were acoustically dispensed into assay ready plates (ARPs) at 3 point 1:5 titrations. ARPs were stored at −20°C and shipped to BSL3 facility (Southern Research Institute, Birmingham, AL) for CPE assay. ARPs were brought to room temperature and 5μl of assay media was dispensed to all wells. The plates were transported into the BSL-3 facility were a 25 μL aliquot of virus inoculated cells (4000 Vero E6 cells/well) was added to each well in columns 3-24. The wells in columns 23-24 contained virus infected cells only (no compound treatment). A 25 μL aliquot of uninfected cells was added to columns 1-2 of each plate for the cell only (no virus) controls.

After incubating plates at 37°C with 5% CO_2_ and 90% humidity for 72 h, 30 μL of Cell Titer-Glo (Promega, Madison, WI) was added to each well. Following incubation at room temperature for 10 minutes the plates were sealed with a clear cover, surface decontaminated, and luminescence was read using a Perkin Elmer Envision (Waltham, MA) plate reader to measure cell viability.

### NS1 TR-FRET assay

HEK293 cells were maintained in EMEM medium with 10% fetal bovine serum, 1% pen/strep (Gibco, Cat. # 15140–122). Cells were seeded at 1000 cells/3 μL/well in the white 1536-well plate and incubated at 37⍰°C with 5% CO2 overnight. Compounds in dilution were added to cells at 23 nL/well and incubated for one⍰hour followed by addition of 2 μL/well of the prototypic ZIKV strain, MR766 solution to cells (MOI⍰=⍰0.5). After an incubation at 37⍰°C for 24⍰h, 2.5 μL/well of detection reagent mixture of two labeled anti-ZIKV NS1 antibodies was added to assay plates. TR-FRET signals were measured using an Envision plate reader (PerkinElmer). Data were normalized by using the control wells (without addition of ZIKV) as a negative control (0% NS1) and positive wells (with ZIKV) as 100% NS1 level.

### In vitro assay and structure data

qHTS data generated on the NPC from the CPE assay (https://opendata.ncats.nih.gov/covid19/index.html) as well as compounds reported as active from recent anti-SARS-CoV-2 repurposing screens^29–31^ and drugs proposed by the scientific community as potential COVID-19 therapies^11,32–34^ were used to train the SARS-CoV-2 models. The detailed qHTS data analysis process including data normalization, correction, classification of concentration response curves, and activity assignment was described previously ^35^. From the CPE assay, compounds that showed concentration dependent response with >30% efficacy were considered active. Other compounds were considered inactive. Literature reported anti-SARS-CoV-2 compounds were considered active.

qHTS data generated in-house at NCATS were used to train the models for ZIKV NS1. NS1 activity data^26^ were generated in qHTS format on three bioactive collections: the Library of Pharmacologically Active Compounds (LOPAC, 1,280 compounds), the NCATS Pharmaceutical Collection (NPC, 2,816 approved and investigational drugs) ^25^, and the Mechanism Interrogation PlatE (MIPE, 1,866 cancer drugs with known mechanism of action) ^36^. Compounds that showed inhibition in both the ratio and 615 nm readouts were considered active. Compounds that were inactive in the ratio readout were considered inactive. Other compounds were considered inconclusive and excluded from modeling. A NCATS in-house collection, the Genesis library, of ~90K diverse compounds was also screened for NS1 activity at a single concentration (14 μM). From these results, compounds that showed >30% inhibition in both the ratio and 615 nm readouts were considered active and other compounds were considered inactive.

The activity data on ~2,600 drugs screened in an EBOV assay from a literature report were used to train the EBOV activity models.^24^ These compounds were mapped to 2,065 unique compounds in the NCATS compound library. The anti-EBOV activities (active or inactive) of these compounds were assigned according to the literature report.^24^

A subset of the compounds in the bioactive collections, NPC and LOPAC in particular, were screened in nearly all the assays available at NCATS. Two NCATS in-house diverse compound libraries, Sytravon, which contains ~44,000 compounds, and Genesis, which contains ~90,000 compounds, and a subset (~100,000 compounds) of the other NCATS bioactive libraries and a large diverse compound library (MLS), were also screened in subpanels of the NCATS assay portfolio. The bioactive compound activity profiles in the assays that also screened the Sytravon (130 readouts), Genesis library (39 readouts), or MLS (225 readouts) were used to train and test the activity-based models (BABM-S or BABM-G). Structure fingerprints were generated for all compounds using the ChemoTyper^37^ for the structure-based models (SBM). Structure data on all the compounds with target activity data available were used to train and test the SBM. The compositions of these datasets are summarized in **Supplementary Table 1** and the different types of models based on these datasets are summarized in **Table 1**. The assay activity-based models (BABM-S, BABM-G, BABM-M) and the activity-structure combined models (CM-S, CM-G, CM-M) were applied to predict the target activity of the compounds with activity profiles available from the Sytravon/Genesis/MLS assays. The SBM was applied to predict the target activity of all ~600K compounds in the NCATS compound library. For activity-based models, only compounds that showed activity in at least 10% of the Sytravon, Genesis or MLS assay panel were kept for analyses. Here, the definition of “active” is not as strict as what would normally be considered as a “hit” for lead identification. Any type of concentration dependent activity observed, regardless of potency or efficacy, was labeled as “active”. As such, compounds that showed activities in multiple assays are not compounds that deemed “promiscuous” in the traditional sense.

**Table 1.** Models built on different training datasets

| Assay Data<br>Source/Model Type | Chemical Structure | Assay Activity | Activity and Structure<br>Combined |
| --- | --- | --- | --- |
| MLS | SBM | BABM-M | CM-M |
| Sytravon | SBM | BABM-S | CM-S |
| Genesis | SBM | BABM-G | CM-G |

SBM = Structure based model
BABM-M = Activity-based model (MLS)
BABM-S = Activity-based model (Sytravon)
BABM-G = Activity-based model (Genesis)
CM-M = Combined model (SBM+ABM-M)
CM-S = Combined model (SBM+ABM-S)
CM-G = Combined model (SBM+ABM-G)

### Modeling

The Weighted Feature Significance (WFS) method previously developed at NCATS^38^ was applied to construct the models. Briefly, WFS is a two-step scoring algorithm. In the first step, a Fisher’s exact test is used to determine the significance of enrichment for each feature in the active compounds compared to inactive compounds, and a p-value is calculated for all the features present in the data set.

For structure data, the feature value was set to 1 for compounds containing that structural feature and 0 for compounds that do not have that feature. For assay activity data, each assay readout was treated as a feature and the feature value was set to 1 for “active” compounds and 0 for inactive compounds. If a feature is less frequent in the active compound set than the inactive compound set, then its p-value is set to 1. These p-values form what we call a “comprehensive” feature fingerprint, which is then used to score each compound for its active potential according to Equation (1), where *p*_*i*_ is the p-value for feature *i*; C is the set of all features present in a compound; M is the set of features encoded in the “comprehensive” feature fingerprint (i.e., features present in at least one active compound); N is the number of features; and α is the weighting factor, which is set to 1 in all the models described here so that all assay features and structure features are treated equally. A high WFS score indicates a strong potential to be active.

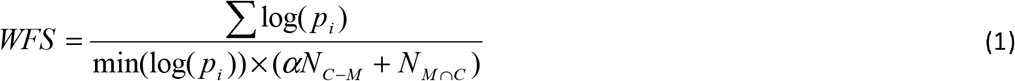

For each model, compounds were randomly split into two groups of approximately equal sizes, one used for training and the other for testing. The randomization was conducted 10 times to generate 10 different training and test sets to evaluate the robustness of the models. Model performance was assessed by calculating the area under the receiver operating characteristic (ROC) curve (AUC-ROC), which is a plot of sensitivity [TP/(TP+FN)] versus (1-specificity [TN/(TN+FP)]) ^39^. A perfect model would have an AUC-ROC of 1 whereas an AUC-ROC of 0.5 indicates a random classifier. The random data split and model training and testing were repeated ten times, and the average AUC-ROC values were calculated for each model. For external experimental validation of models, model performance was measured by the positive predictive value (PPV = TP/(TP+FP)). Statistical significance was determined by the Fisher’s exact test comparing model PPV with the active rate in the training dataset for the corresponding target being modeled.

### Selection of model predicted actives

Models with AUC-ROC >0.75 were considered for compound selection. WFS score cutoff values for model predicted actives were determined using the ROC curves where both sensitivity and specificity were optimized. Only compounds that scored higher than the cutoff values were considered candidates for follow up selection. Due to the limitations of different assays and resources, for each target we selected compounds with the largest possible structure diversity that could fit into one 1,536-well plate for experimental validation. When the candidate pool was much larger than the target number of compounds, the candidates were narrowed down based on structure type. For this purpose, the entire NCATS in-house compound library was clustered based on structure similarity (729-bit ChemoTyper^37^ fingerprints) using the self-organizing map (SOM) algorithm^40^. From the clusters that contain model predicted actives, a fraction of the active compounds was selected from each cluster based on the WFS score and the number of models that predicted the compound as active. Because the EBOV assay could only test ~100 compounds, the anti-EBOV candidates were manually inspected and narrowed down further based on literature reports, structure novelty and ADME properties. In most cases the selection was driven by availability of physical samples. All compounds that met the WFS score cutoff from a model were selected when less than 1,408 compounds had physical samples available for cherry picking. The SARS-CoV-2 CPE assay (live virus) could only be run in 384-well format. Limited by the testing space available and physical sample availability, only 311 model predicted compounds were selected for experimental confirmation in the SARS-CoV-2 live virus assay.

## Results

### Model performance and validation

**Table 2** provides an overview of the three viral targets (SARS-CoV-2, ZIKV, EBOV) used for modeling. The entire model training, testing and validation process is illustrated in **Figure 1**. Model performance was measured by the area under the ROC curve (AUC-ROC; see Methods section for details). The majority of the models performed well on their corresponding test sets with mean AUC-ROC values >0.8 (**Figure 2A** and **Supplementary Table 2**). The structure-activity combined models (CM) showed the best performances compared to the models built on activity (BABM) or structure (SBM) alone with mean AUC-ROC values >0.83. Of the BABM models using data from different assay panels and compound libraries, the BABM-S and BABM-M models all showed good performances with mean AUC-ROC values of 0.79 and 0.84, respectively (**Supplementary Table 2**). The BABM-G models with the smallest assay panel for training showed the lowest AUC-ROC values averaging 0.75. The structure-based models generally showed lower performances than the CM and BABM models with a mean AUC-ROC of 0.72. **Supplementary Figure 1** shows example ROC curves from each of the three types of models.

**Figure 1.**
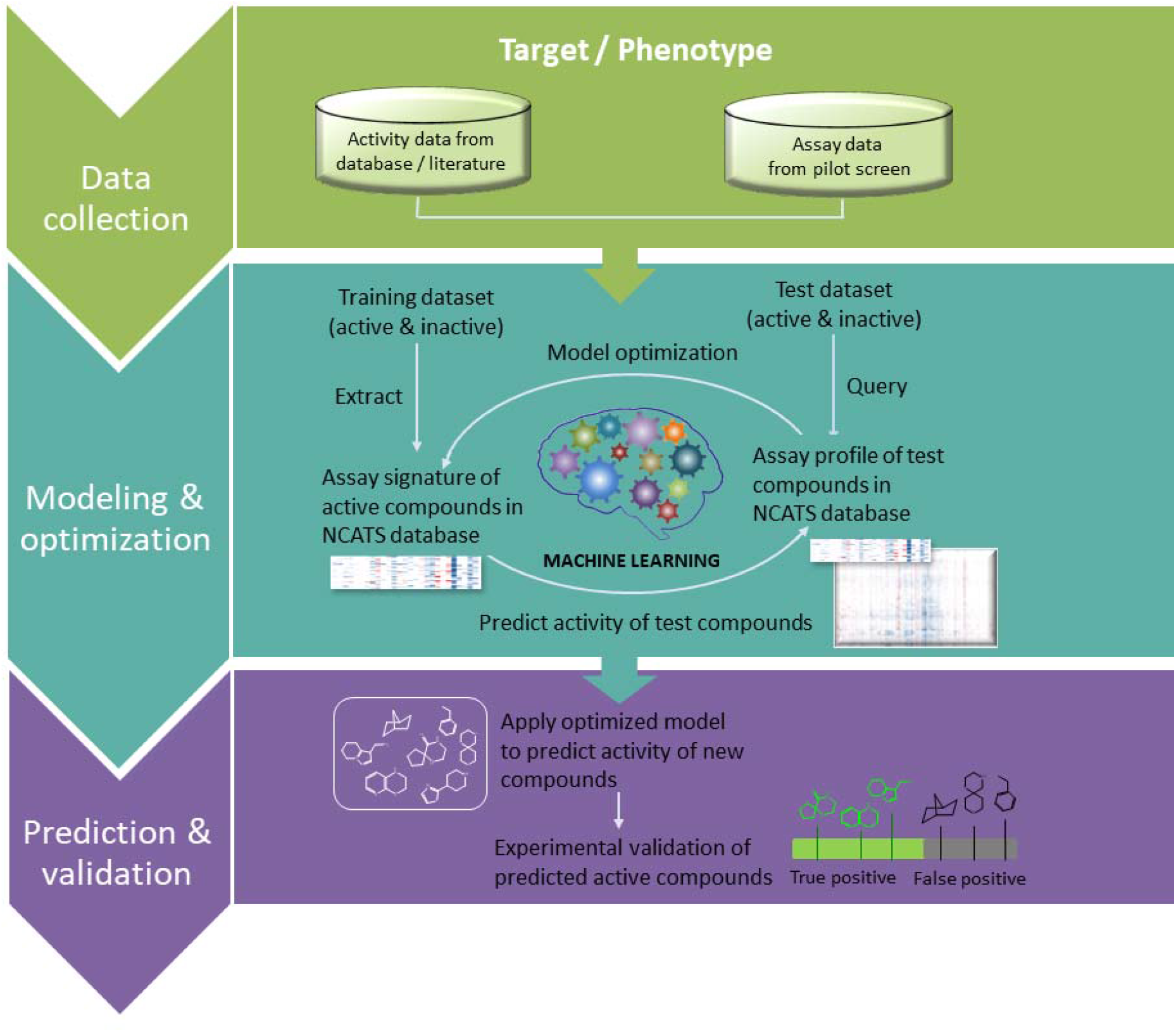
Biological activity-based modeling (BABM) process. For any biological target of interest, *T* (e.g., SARS-CoV-2, ZIKV NS1, EBOV), the model identifies the activity pattern of active vs. inactive compounds based on the training data, which are activity profiles of a set of compounds across a diverse panel of assays including *T*. The active signature is then matched against the activity profiles of a new set of compounds across the same assay panel. The ability of the model to use this signature to correctly identify actives from the new compound set is first tested using part of the data with known *T* activity (the test set). An AUC-ROC value is calculated using the test set to evaluate the model performance. The model is then applied to a set of compounds with unknown *T* activity (prediction set; e.g., Sytravon, MLS, Genesis). Predictions are made on the new compounds based on their activity profile similarity to that of the active signature for *T*. The predicted *T* actives are further validated experimentally for their activity against *T*. Comparing experimental results with model predictions, true positives (TP) and false positives (FP) are counted to determine the performance of the model. In the heat maps, each row represents a compound, each column is an assay, and the heat map is colored by the compound activity.

**Figure 2.**
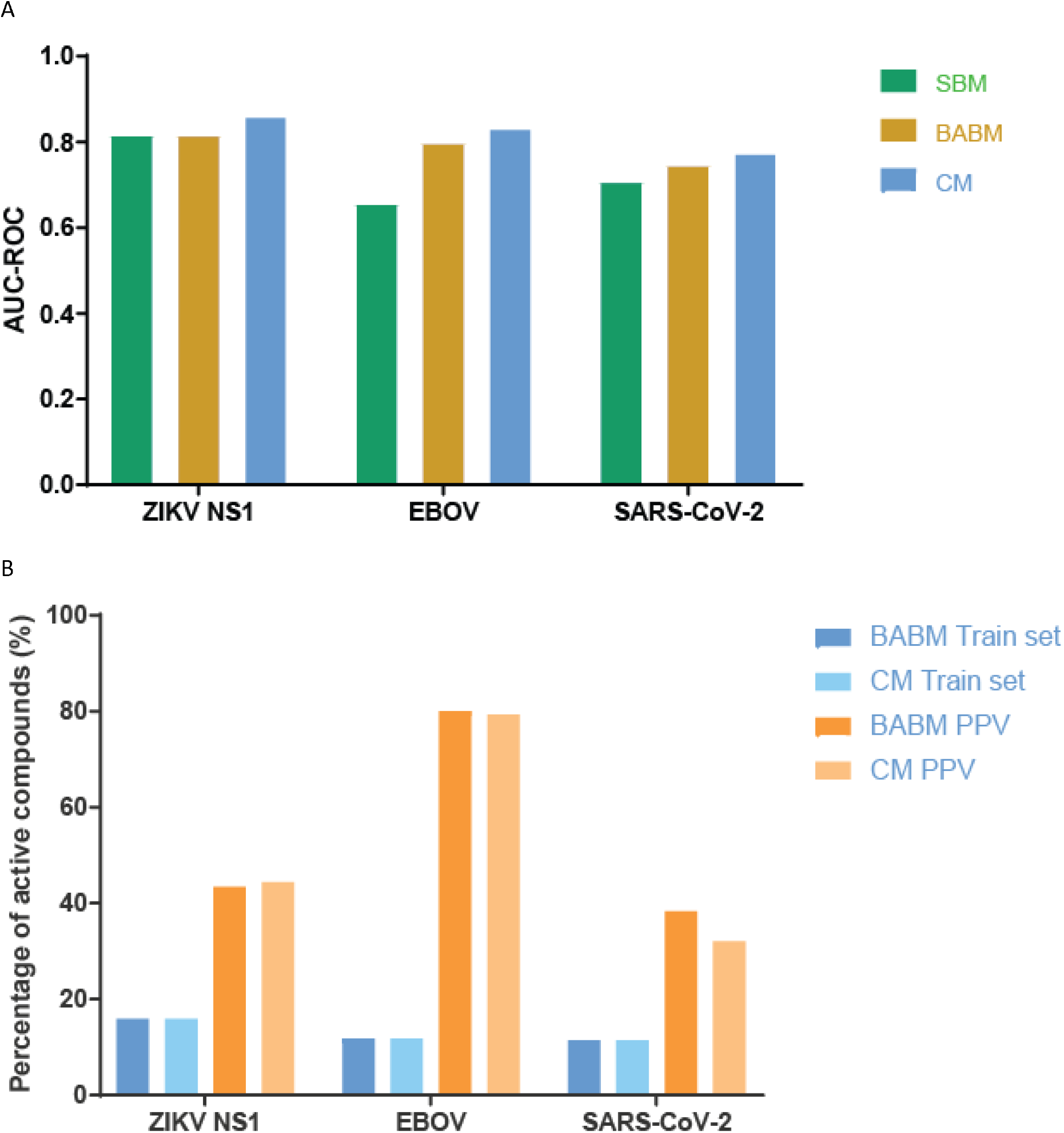
Model performance and experimental validation. A. Model performances on the test sets measured by AUC-ROC values. B. Model performances measured by external experimental validation PPV (colored in different shades of brown) in comparison to training set active rates (e.g., original assay hit rate; colored in different shades of blue). Model selected compounds are significantly enriched with true actives. Model type: SBM = Structure based model; BABM = Activity-based model (Sytravon); CM = Combined model (SBM+BABM).

**Table 2.** Overview of viral targets used for modeling and summary statistics of model identified active compounds.

| Model | Target | Data Source | Data Type | Active Enrichment<br>(fold)* | Potency Range** | Structure Diversity*** |
| --- | --- | --- | --- | --- | --- | --- |
| **SARS-CoV-2** | No | In house/Literature | HTS/Literature | 2.8-3.3 | <1 μM - 32 μM | 0.07-0.23 |
| **ZIKV NS1** | Yes<sup>a</sup> | In house | qHTS | 2.7-27.5 | 1 nM - 63 μM | 0.04-0.26 |
| **EBOV** | No | Literature | HTS | 6.5-6.8 | 0.4 μM - 25 μM | 0.07-0.21 |

<sup>a</sup> The NS1 protein is a nonstructural protein that is not present in the virus itself but only appears in the host cells when virus starts replication. The NS1 assay detects compounds that block NS1 protein production, which may inhibit virus entry to cells, and virus RNA and virus protein productions in cells.
\* Fold = hit rate of model predicted actives/hit rate of assay used for model training
\*\* Potency range of experimentally confirmed model predicted actives
\*\*\* Tanimoto similarity was calculated between each model predicted active and active compounds in the training set. The values shown here are the range of the average Tanimoto scores for the model predicted actives.

To further validate the models and identify new compounds with antiviral activity, a subset of model predicted actives was selected for each viral target for experimental validation. For ZIKV, 1,676 selected actives predicted by the model were tested in the original NS1 assay^26^ that generated the data to train the models. To validate the EBOV models, the EBOV-eGFP infection assay^26,41^ was applied to test 96 selected model predicted EBOV actives. All 96 compounds were first inspected at 30 μM for potential cytotoxicity, resulting in 62 compounds with <50% cell killing, which were further tested for EBOV infection inhibition. The EBOV inhibition activity of these 62 compounds were used to evaluate model performance. The positive predictive value [PPV = TP/(TP+FP)], i.e., the fraction of model predicted actives that are experimentally confirmed, was calculated for each model (**Figure 2B** and **Supplementary Table 2**). The model PPVs ranged from 30% (SBM for NS1) to 89% (CM-G for EBOV). The EBOV models showed higher PPVs (~80%) than the NS1 models (~40%). Compared to the active rates in their corresponding training datasets (i.e. original assay hit rate), all model predicted active sets were significantly enriched with true active compounds (Fisher’s exact test: p<10^−10^) (**Supplementary Table 2**). For example, the active rate of the EBOV BABM-S model training set was 11.8% and the corresponding model PPV (i.e. experimental validation set active rate) was 80%. Thus the enrichment of actives by the EBOV BABM-S model was 6.8-fold (80/11.8). The enrichment of actives for all models (**Table 2**) ranged from 2.7-fold (BABM-S for NS1; p<10^−20^) to 27.5-fold (SBM for NS1; p<10^−20^). Most models showed enrichments between 5- and 10-fold when compared to the active rates in the training set. The potency ranges of the experimentally confirmed actives are summarized in **Table 2** and **Figure 3**. The models identified potent compounds with novel structures for all four disease targets with IC_50_s in the nanomolar range (Figure 3). Experimental validation data for all models are provided as **Supplementary Data 1**.

**Figure 3.**
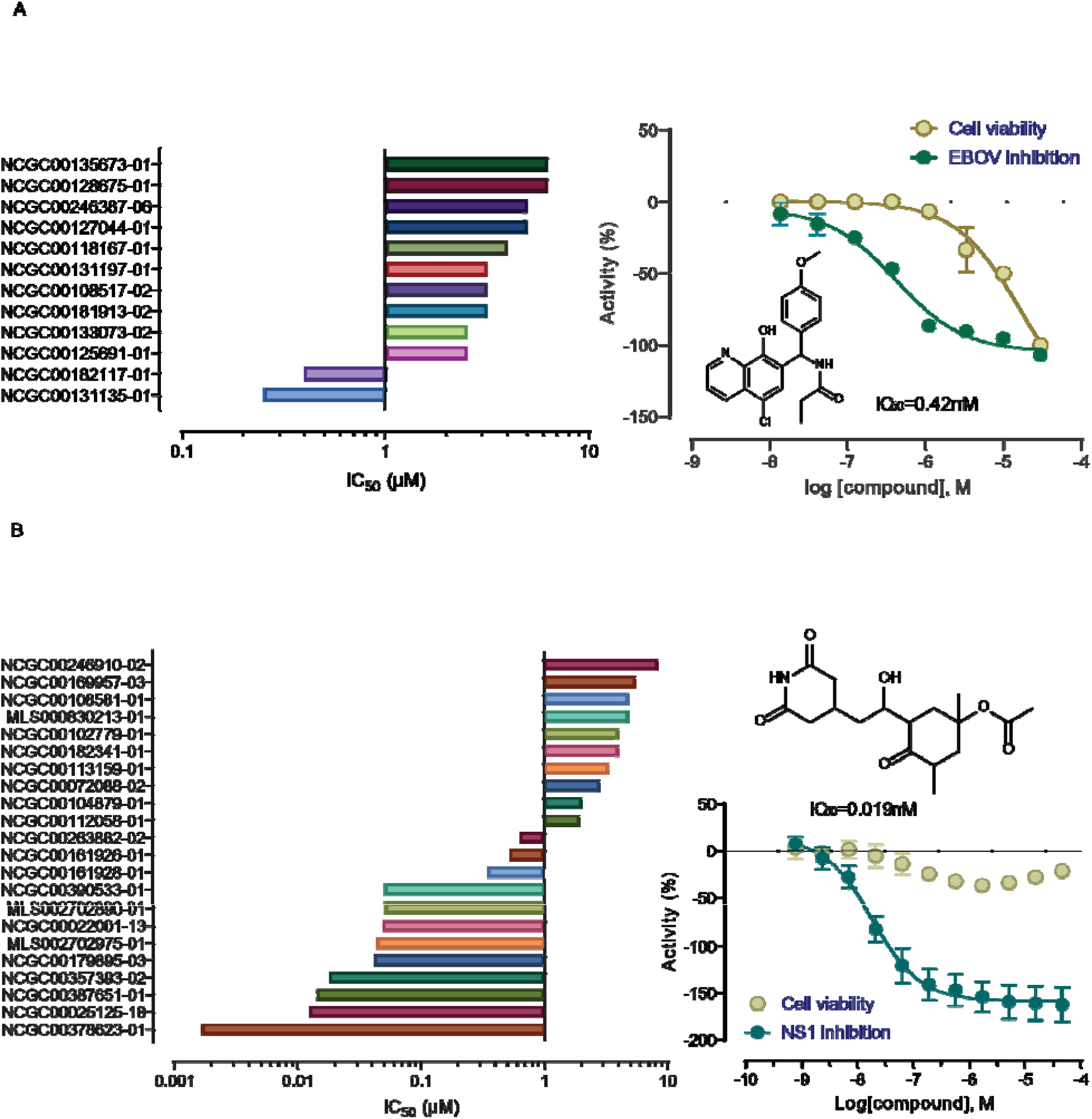
Experimental validation results of model predicted actives. A. Potencies and examples of compounds confirmed in the EBOV inhibition assay with minimal cytotoxicity. B. Potencies and examples of compounds confirmed in the ZIKV NS1 inhibition assay with minimal cytotoxicity.

### Identification of novel anti-EBOV and anti-ZIKV compounds

Of the 50 compounds with anti-EBOV activity confirmed at 30 μM, we selected 27 that showed >90% inhibition of EBOV infection with minimal cytotoxicity (>80% cell viability) to test in concentration-response format (0.17 nM to 30 μM; 1:3 fold dilution; triplicate) to determine their EBOV inhibition potency. All 27 compounds showed concentration dependent inhibition of EBOV infection with IC_50_s ranging from 25 nM to 25 μM (**Figure 3A**; **Supplementary Data 2**). Seven of these compounds were potent with IC_50_≤5 μM and were not apparently cytotoxic or at least six times more potent in the EBOV inhibition assay compared to the cell viability counter assay. Two of the seven compounds are known drugs. Arbidol (or umifenovir) (IC_50_~5 μM) is an antiviral treatment for influenza infection used in Russia and China.^42^ A more recent study reported *in vitro* activity of umifenovir at preventing entry of the EBOV Zaïre Kikwit.^43^ Difeterol is used as an antihistamine drug in Japan.^44^ Difeterol (IC_50_ ~3 μM) was also identified as one of the active compounds from a high throughput screen using the EBOV entry assay.^23^ The other five compounds have novel structures with no previously reported anti-EBOV activity. These compounds have the potential to be developed into new antiviral therapies.

A subset (170) of the experimentally confirmed NS1-assay active compounds with relatively potent NS1 signal inhibition activity (IC_50_ <10 μM) and no apparent cytotoxicity were selected for secondary confirmation with compounds tested at 11 concentrations in triplicate (**Figure 3B**; **Supplementary Data 2**). Ten of the 170 compounds did not show activity in the secondary confirmation assay yielding a confirmation rate of 94% for the NS1 assay. 29 compounds showed potent inhibition with IC_50_ <1 μM, 17 of which were not apparently cytotoxic or at least three times more potent in the NS1 assay. A number of these potent compounds are known drugs or bioactives. Colchicine, podofilox, 7-epi-docetaxel and β-peltatin are all microtubule inhibitors. Colchicine is a medication most commonly used to treat gout and as an anti-inflammatory drug for a number of other conditions. Podofilox, or podophyllotoxin, is used as a medical cream to treat genital warts and molluscum contagiosum. 7-Epi-docetaxel is an impurity of docetaxel, a chemotherapy medication used to treat a number of types of cancer. β-Peltatin is a plant metabolite with antineoplastic properties that belongs to the same structural class as podophyllotoxin.^45^ Dolastatin 10, a pentapeptide, is also an inhibitor of microtubule assembly and an investigational drug that has been in clinical trials for its antineoplastic activity.^45^ Like colchicine, narciclasine, an amaryllidaceae alkaloid, was intensively investigated as an antitumor compound both *in vitro* and *in vivo* and have shown anti-inflammatory actions *in vivo*.^46^ Floxuridine is a nucleoside that belongs to the class of antimetabolites. Floxuridine is an oncology drug most often used in the treatment of colorectal cancer. The other known bioactive compounds include two mycotoxins, diacetoxyscirpenol and T-2 Toxin. The other eight potent compounds have novel structures that can potentially be developed into new antiviral therapies.

### Identification of novel anti-SARS-CoV-2 compounds

The activity of 311 compounds predicted by the SARS-CoV-2 BABM models were tested in the live virus CPE assay, 99 of which were confirmed as active, yielding a hit rate of 32% (**Figure 2B** and **Supplementary Table 2**). The model PPVs ranged from 32% (CM-S) to 38% (BABM-S). Compared to the active rates in their corresponding training datasets, all model predicted active sets were significantly enriched with true active compounds (Fisher’s exact test: p<10^−3^) (**Supplementary Table 2**). Compared to the hit rate of the original NPC screen (11%), the models were able to improve the hit rate by 2.8- to 3.3-fold (**Table 2**). The SBM was not used for compound selection because its performance (average AUC-ROC = 0.71) during model training and testing did not meet the 0.75 cutoff. Nonetheless, the SBM predictions made on the 311 compounds were used to assess the performance of the SBM on the experiment validation set in comparison with the BABM models (**Figure 2B** and **Supplementary Table 2**). The PPV of the SBM was 31.6%, lowest of all SARS-CoV-2 models. The potency ranges of the experimentally confirmed actives are summarized in **Table 2** and **Figure 3**. Experimental validation data for all 311 compounds are provided in **Supplementary Data 1**. The structures and WFS scores of ~5,000 compounds predicted as active by at least one of the SARS-CoV-2 BABM models are provided as **Supplementary Data 3**.

The experimentally confirmed SARS-CoV-2 active compounds were further tested at 8 concentrations (instead of 3 concentrations in the primary screen) to get more accurate potency measures (**Supplementary Data 2**). Five of the 95 compounds did not show activity in the secondary confirmation assay yielding a confirmation rate of 95% for the SARS-CoV-2 CPE assay. The most potent compound (MLS000699212-03; Benzaldehyde, 3-methyl-, 2-(2,6-di-4-morpholinyl-4-pyrimidinyl)hydrazone) had an IC_50_ of 500 nM. This compound has only one published study, which is a patent on a compound series described as autophagy modulators for treating neurodegenerative diseases.^47^ Autophagy has been implicated in the entry of coronavirus into host cells, including SARS-CoV, MERS-CoV and SARS-CoV-2^48,49^. Another potent compound with IC_50_ <1 μM (800 nM) is a completely novel synthetic compound with no previous literature report (NCGC00100647-01; N2,N4-bis(3-methylphenyl)-6-(4-morpholinyl)-1, 3,5-Triazine-2,4-diamine). In addition, 13 compounds had IC_50_ <5 μM. Spiramide (AMI-193) is an experimental antipsychotic that acts as a selective 5-HT2A, 5-HT1A, and D2 receptor antagonist.^50^ Ftormetazine is a derivative of the phenothiazine class of antipsychotic drugs.^51^ Prenylamine is a calcium channel blocker of the amphetamine chemical class, which was used as a vasodilator in the treatment of angina pectoris and later withdrawn due to cardiotoxicity. ^52,53^ BEPP [1H-benzimidazole1-ethanol,2,3-dihydro-2-imino-a-(phenoxymethyl)-3-(phenylmethyl)-,monohydrochloride] is a synthetic compound that has been reported to inhibit viral replication by inducing RNA-dependent protein kinase-dependent apoptosis.^54^ Triparanol was the first synthetic cholesterol-lowering drug which was introduced in the U.S. in 1960, but was later withdrawn due to severe adverse effects.^55,56^ Octoclothepine is a tricyclic antipsychotic drug for the treatment of schizophrenic psychosis with high affinities for the dopamine receptors.^57^ Metaphit, the m-isothiocyanate derivative of phencyclidine, is an investigational drug that acts as an acylator of NMDARAn, sigma and DAT binding sites in the CNS.^58^ L-703, 606 is a known antagonist of the neurokinin-1 (NK1) receptor.^59^ The other 5 are novel compounds without any well annotated biological activity.

## Discussion

Traditional QSAR models rely on chemical structure similarity to infer biological activity and thus are limited in their power to discover new chemical scaffolds. Consequently biological activity predictions made on chemicals with structure types not included in the training set are often not reliable – this is commonly referred to as the “applicability domain” (AD) issue.^60^ QSAR models are thus fundamentally restricted by their ADs, namely by the chemical spaces within which the models were originally trained. Incorporating biological response patterns into the models helps to alleviate this issue by expanding the model AD to cover structurally dissimilar chemicals that share similar activity profiles. Activity-based modeling is a relatively new concept, especially when applied to drug discovery. The prerequisite of activity-based modeling is the availability of sets of compounds tested consistently across multiple biological assays with the results serving as compound descriptors or fingerprints. This is enabled by the recent advances in HTS technologies that have produced a tremendous amount of biological activity data in a relatively short amount of time. As a center specialized in HTS, NCATS has a data repository that hosts biological response data on over half a million compounds tested against thousands of assays mostly in qHTS format, which form a rich set of activity profiles at unprecedented scale (over 130 million wells screened over the last 4 years).^3,61^ We show here that a subset of these data could be used to build activity-based models to identify novel antiviral compounds for Zika and Ebola. After validating the robustness of these models, we applied the BABM approach to build predictive models for the novel SARS-CoV-2 producing lead compounds, confirmed in a live virus assays, with potential of being developed into new COVID-19 therapies.

BABM was applied to ZIKV NS1 models as a proof-of-concept comparison to traditional modeling approaches. **Supplementary Figure 2** shows the 652 experimentally confirmed NS1-expression actives correctly identified by at least one of the three models (SBM, BABM, and CM). Only 43 of the 652 compounds were predicted as active by all three models. BABM identified 82%, while the CM and SBM identified 76% and 28% of 652 active compounds, respectively. BABM and CM identified the majority of the actives, most of which (448 out of 534) were also shared by both models. SBM identified a much smaller number of actives, most of which (118 out of 182) were also different from those identified by the BABM and CM. Out of the 118 actives picked up by the SBM only, 106 were not predicted by the BABM or CM because no assay activity profile data were available on these compounds. As structure-based models rely on structure similarity to make predictions, these models were not reliable in predicting compounds with completely new scaffolds that were not already represented in the training set. Activity-based models which rely on activity profile data for training, in contrast, are not restricted by structure similarity and thus are potentially more powerful in discovering new scaffolds. Models for all three viral targets in this study identified compounds with very low structural similarity to active compounds in the training set (e.g., average Tanimoto similarity scores to the training set as low as 0.04-0.07; **Table 2**). Compared to traditional QSAR models built with chemical structure data alone, the BABM identified compounds that are structurally distinct from the training set and the compounds identified by the SBM (**Figure 4**), demonstrating the advantage of the BABM in discovering new chemical types.

**Figure 4.**
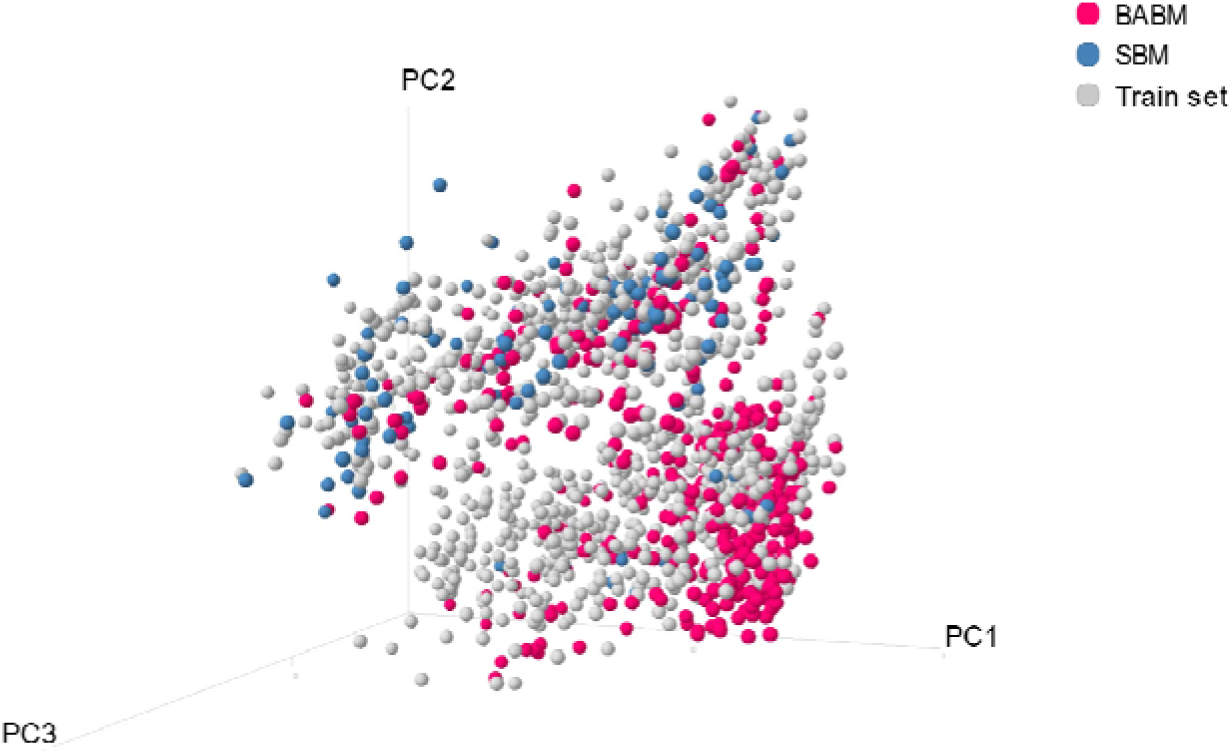
Chemical structure space occupied by compounds predicted as active by the BABM and SBM NS1 models, and active compounds in the training set, i.e., the original NS1 screen. The plot shows that the compounds identified by the activity-based model occupied structure spaces distinct from those identified by the structure-based model.

Combining traditional structure-based models with BABM can maximize the chance of identifying the best novel lead compounds as new candidates for any therapeutic target of interest. Both the BABM and CM models used activity data in other assays as descriptors for training while the CM used structure features in addition. The model predictions were further validated experimentally. Again using the NS1 models for example, even though the BABM identified a larger portion of the experimentally confirmed actives (i.e., was more sensitive), the CM had a lower false positive rate (i.e., was more specific). Adding structure information helped the CM to achieve a slightly improved PPV. For all three viral targets modeled in this study, the CM models achieved the best overall performance compared to the SBM and BABM models. More intriguingly, the sizes of the training sets for all the models were much smaller than the prediction sets on which the models were applied, 30- to 100-fold for the BABM and CM models, and up to 300-fold for the SBM (**Supplementary Table 1**). That models built on a small training set performed well on predicting a much larger and more diverse set of compounds with accuracies on par with or better than most *in silico* screening approaches further demonstrated that the models were robust enough to be applicable to large and diverse compound collections to identify new leads.^1,62^

The SARS-CoV-2 BABM models identified ~100 compounds that were experimentally verified to show antiviral activity in a live virus assay. Some of these compounds have novel structures that could be developed into new classes of antiviral therapeutics. Models built for Zika and Ebola also identified potent new lead compounds. In addition, we provided the prediction results of ~5,000 compounds that were predicted as active by the SARS-CoV-2 BABM models as a resource to the scientific community to develop new anti-COVID-19 therapies. The activity-based approach was demonstrated here to be able to be rapidly applied to identify new lead compounds for novel targets or disease phenotypes.

As a complement to structure-based approaches, either ligand or target structure-based, the additional information provided by activity data is shown here to significantly improve the predictive power of VS models. The novel chemical scaffolds identified by BABM from an existing screening library can also be incorporated into new QSAR models to screen other chemical libraries more efficiently with no bioactivity profiles available. Of note is that, in addition to HTS libraries, the general concept of BABM can be extended to any type of biological data, such as genomics and proteomics data, data generated on mixtures or antibodies, and clinical data, where clearly defined structure information is not available. As such the BABM approach shows the promise of broad applications in different areas of biology.

## Acknowledgements

This work was supported by the Intramural Research Programs of the National Center for Advancing Translational Sciences, National Institutes of Health. The authors would like to thank Hui Guo, Xin Hu, and Min Shen for assistance with CPE assay data processing, and Richard Eastman, Zina Itkin, and Paul Shinn for compound management and plating.

## Author Contributions

R.H., W.Z., W.H. and C.P.A. conceived the research and designed the study. M.X., H.Z., C.Z.C., E.M.L. and S.H. performed the experiments, collected data and aided data interpretation. R.H. performed modeling and statistical analysis of all data. H.Z. aided data analysis and visualization. K.S. and D.B. aided compound selection. R.H. and W.Z. wrote the manuscript. R.H., W.H., M.D.H., D.L., A.S., C.P.A., X.Q., H.T. and W.Z. directed the research. All authors reviewed the manuscript.

