## Supplementary Information for "Massive-scale biological activity-based modeling identifies novel antiviral leads against SARS-CoV-2"

**Table S1**. Training dataset compositions for models

| SARS-CoV2 Data Set | Feature Count | Active Count | Inactive Count | Prediction Set |
| --- | --- | --- | --- | --- |
| SBM | 729 | 279 | 2155 | 617,947 |
| BABM-S | 130 | 209 | 860 | 109,291 |
| BABM-M | 225 | 200 | 940 | 99,660 |
| CM-S | 859 | 209 | 859 | 103,400 |
| CM-M | 954 | 200 | 939 | 96,142 |

| NS1 Data Set | Feature Count | Active Count | Inactive Count | Prediction Set |
| --- | --- | --- | --- | --- |
| SBM | 729 | 1,023 | 91,643 | 527,715 |
| BABM-S | 130 | 281 | 1,486 | 108,593 |
| CM-S | 859 | 281 | 1,486 | 102,701 |

| EBOV Data Set | Feature Count | Active Count | Inactive Count | Prediction Set |
| --- | --- | --- | --- | --- |
| SBM | 729 | 148 | 1,833 | 618,400 |
| BABM-S | 130 | 128 | 958 | 109,274 |
| BABM-G | 29 | 110 | 710 | 70,932 |
| CM-S | 859 | 128 | 956 | 103,384 |
| CM-G | 758 | 108 | 704 | 64,397 |

**Table S2**. Model performances

| SARS-CoV2 Model | AUC-ROC (Test Set) | Cherry Pick Validation | | | Training Set Active Rate* | P |
| --- | --- | --- | --- | --- | --- | --- |
|  |  | TP | FP | PPV |  |  |
| SBM** | 0.71±0.01 | 12 | 26 | 31.58% | 11.46% | 8.97×10^-4^ |
| BABM-S | 0.75±0.02 | 28 | 45 | 38.36% | 19.55% | 4.64×10^-4^ |
| BABM-M | 0.79±0.02 | 25 | 44 | 36.23% | 17.54% | 3.47×10^-4^ |
| CM-S | 0.77±0.02 | 83 | 176 | 32.05% | 19.57% | 2.66×10^-5^ |
| CM-M | 0.81±0.02 | 60 | 100 | 37.50% | 17.56% | 3.84×10^-8^ |

*The training set active rate for the SBM is close to the SARS-CoV2 CPE qHTS assay hit rate. The training set active rates for the activity-based models are higher because the NPC screened for this assay was recently updated with many new drugs not in the older version. Profile data in other assays were not available for these new drugs, most of which were inactive in the CPE assay, thus they were not included in training the activity-based models.

**Not used to select compounds for experimental validation.

| NS1 Model | AUC-ROC (Test Set) | Cherry Pick Validation | | | Training Set Active Rate | P |
| --- | --- | --- | --- | --- | --- | --- |
|  |  | TP | FP | PPV |  |  |
| SBM | 0.82±0.01 | 182 | 419 | 30.28% | 1.10% | <10^-20^ |
| BABM-S | 0.82±0.02 | 532 | 692 | 43.46% | 15.90% | <10^-20^ |
| CM-S | 0.86±0.01 | 493 | 617 | 44.41% | 15.90% | <10^-20^ |

| EBOV Model | AUC-ROC (Test Set) | Cherry Pick Validation* | | | Training Set Active Rate | P |
| --- | --- | --- | --- | --- | --- | --- |
|  |  | TP | FP | PPV |  |  |
| SBM | 0.66±0.02 | N/A | N/A | N/A | 7.47% | N/A |
| BABM-S | 0.80±0.02 | 48 | 12 | 80.00% | 11.79% | <10^-20^ |
| BABM-G | 0.70±0.02 | 13 | 2 | 86.67% | 13.41% | 7.05×10^-10^ |
| CM-S | 0.83±0.01 | 46 | 12 | 79.31% | 11.81% | <10^-20^ |
| CM-G | 0.78±0.03 | 16 | 2 | 88.89% | 13.30% | 3.04×10^-12^ |

*Cherry picked 96 compounds, 34 of which killed cells at 30 µM and not tested for Ebola. The remaining 62 compounds were used to evaluate model performance.

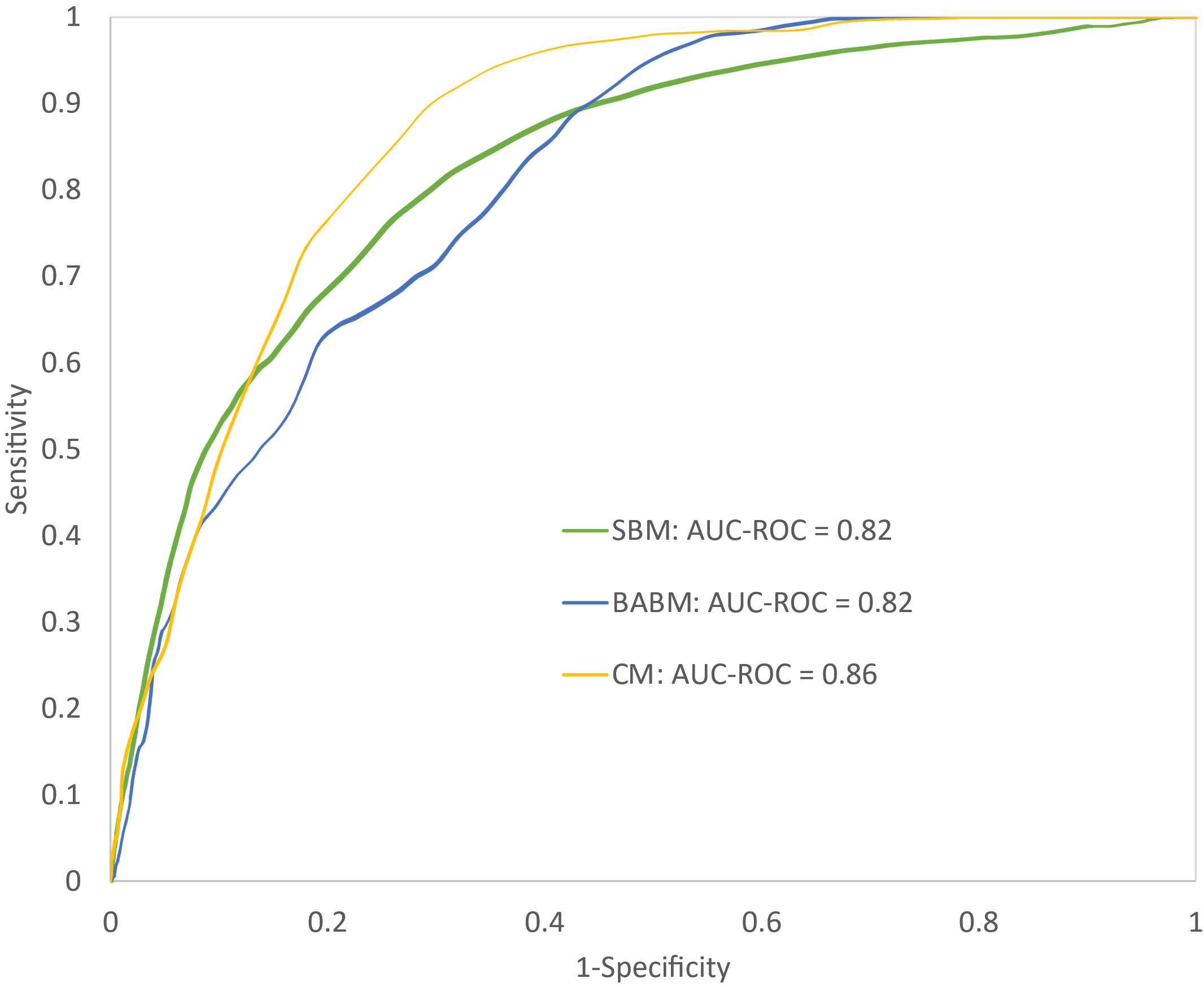

**Figure S1**. Example ROC curves on the test set. Curves were selected from the NS1 models. BABM = biological activity-based model; SBM = structure-based model; CM = combined model.

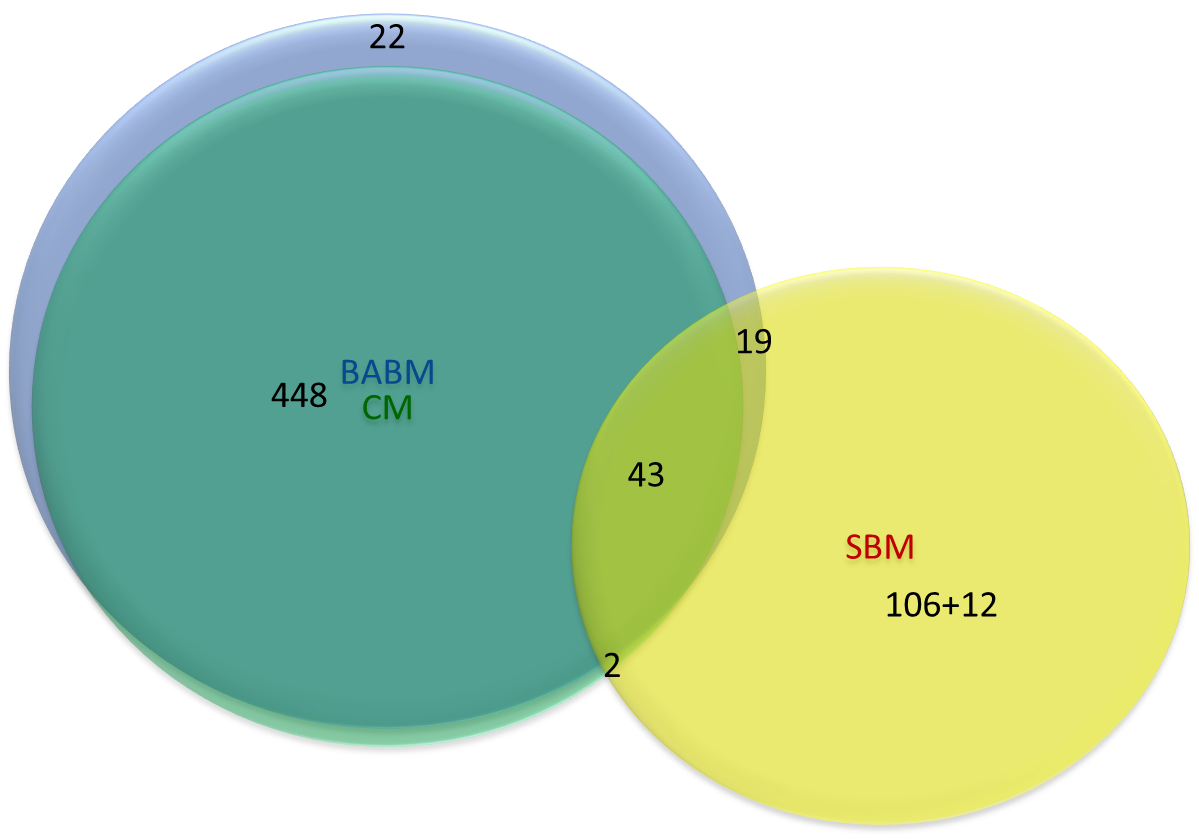

**Figure S2**. Confirmed NS1 actives correctly identified by each model. BABM = biological activity-based model; SBM = structure-based model; CM = combined model.
